# Remote inferences and direct observations provide complementary insights into foraging behavior

**DOI:** 10.1101/2025.03.05.641645

**Authors:** Jack Graham Hendrix, Eric Vander Wal

## Abstract

Behaviorists sometimes view askance studies where researchers indirectly observe animals, consequently challenging whether remotely inferred behavior is true behavioral research. Alternatively, others purport that technological advancements, like Global Position System (GPS) tags or biologgers, have expanded the scope of behavioral research to temporal and spatial scales infeasible for direct observation. To spotlight strengths and shortcomings in approaches to behavioral research, we interrogated the use of techniques and their assumptions in foraging research, a behavior of interest to ecologists and behaviorists. We reviewed 604 foraging behavior studies to synthesize and compare foraging research across disciplines, taxa, and methods. We sorted approaches by the data they collect and their associated assumptions and determined that rather than two categories of direct vs. remote, there were five: direct observation, tracking, biologger, remote audio-visual, and remote spatial. Categories differed in their spatial extents, with remote spatial research having a much larger extent (up to 1.6 million km^2^) than direct observational or remote audio-visual studies. Remote spatial studies also spanned large temporal extents, but temporal coverage (the proportion of total study duration when data are actively collected) was lower compared to biologger research. Methods were also applied to different stages of the behavioral process of foraging: direct observations and tracking involved searching for resources at finer scales. 23% of studies used > 2/5 categories. A compound approach provided a more nuanced and complete description of foraging behavior. Thus, our understanding of behavior improves when multiple approaches are applied in conjunction to our question of interest.

**Lay summary:** How do you study behavior? Your first response might be simply watching animals behave. While there are many ways to indirectly infer behavior without directly observing wildlife, some question these approaches because we cannot be 100% confident in the inferred behavior. We reviewed >600 behavior studies and found that though remote inferences rely on assumptions, so do direct observations. Remote inferences also permit studying temporal and spatial scales infeasible when watching wildlife directly.

## Introduction

Fifty years ago, Altmann (1974) published her germinal paper outlining the applications of field observations of animal behavior. Altmann contrasted field studies of behavior with the prevailing practice of observing captive animals, noting that “*one hears the claim that accurate studies of behavior can be made only in the laboratory, and that quantitative research on behavior is not practicable in the context of ongoing, real-life situations.”* Half a century later, direct observations of wildlife in the field have become the foundation of behavioral ecology. Now established as the dominant method, a similar parallel is sometimes drawn today between direct observations and remotely inferred behavior observed indirectly through remote technology.

Methods to study behavior today are often sorted into two main categories based on how the data are collected: direct behavioral observations, grounded in the field of classical ethology (Altmann, 1974), and remotely sensed behavior through biologgers or spatial data (Börger et al., 2020). Direct behavioral observations provide concrete information about how animals are behaving at a specific time and place, while remotely sensed approaches rely on inference and assumption to convert data streams such as Global Position System (GPS) movement paths or accelerometer output into behavioral information (Ropert-Coudert & Wilson, 2005). However, this dichotomy oversimplifies the diversity of approaches. For example, motion-activated video cameras blur the lines between direct and remote methods. Many direct observational studies are also mediated by technology as video recording of study populations has become de rigueur, addressing many of the feasibility concerns of observing all simultaneous behaviors in a group as discussed by Altmann (1974).

Given this complexity, approaches to studying wildlife behavior might be better categorized by the type of information they produce and the assumptions underlying their interpretation, rather than the method or technology used to obtain the data. While remote inferences have limitations and do require assumptions to be made about the relationship between data output and the actual behavior, classical behavioral research also has limitations and important assumptions. Direct observations are limited to situations with sufficient daylight and open habitat to observe individuals, so assumptions must be made when extrapolating behaviors across space and time, but this assumption is rarely acknowledged. Observer effects are more commonly noted as a potential influence on behavior, though habituation protocols can alleviate some of these concerns (Rankin et al. 2009).

To understand better the types of insight that remote behavioral research can provide relative to direct observations, and to clarify the data type resulting from different research methods, we performed a literature review using foraging as our behavior of interest. The history of the study of foraging makes it an apt case study. Foraging generated much early interest from behavioral ecologists because of its downstream implications for life history and population ecology (Lemon 1991). Classical ethology and direct observations of feeding behavior formed the foundation of the study of foraging ecology (Kamil et al. 1987), but foraging is now a topic of interest for many types of ecologists outside of behavioral ecology. The expanded interest in foraging, facilitated by technological advances, brings with it different preconceptions and understandings of foraging as a concept from other subdisciplines, such as spatial ecology. Our working definition of foraging for the purposes of this review is “the seeking out, acquisition, and consumption of food resources” (see Appendix 1 for details).

### Ethology and foraging

Ethology refers to the study of animal behavior and describes the collection of approaches involving direct observations of animals and recording their behavior. The precise methods vary by system and question. Quantitative analyses of behavior typically interrogate the behavioral response of interest either as events to be counted, or states with a duration as part of a time budget (Altmann 1974). Sampling procedures may involve following focal individuals for extended intervals or instantaneous scan sampling of groups at regular points in time. Data collection can include every behavior observed as part of an ethogram that is either generalized to some degree (MacNulty et al. 2007; Stanton et al. 2015) or specific to the study species.

A complication that arises from direct observation is the influence of the observer. It is challenging to directly observe wildlife while ensuring the act of observation does not disturb the wildlife. There is a risk that the behavior observed by researchers does not reflect the animal’s behavior when humans are absent. While it is common practice in observational studies to habituate study subjects to the presence of humans (Johns 1996), and to validate the success of this process (Gazagne et al. 2020),

For cryptic or far-ranging species, records of foraging and feeding behavior are difficult to obtain by direct observation. Even for easily observed and habituated species, the process of observing and recording behavior for hours each day is laborious (Schuppli et al. 2016). With the advent of radio telemetry and satellite technology since the 1970s, a new avenue for foraging research in previously inaccessible situations arose (Tomkiewicz et al. 2010).

### Remotely-sensed foraging data

For the purposes of this review we will be using remote sensing to refer to biotelemetric approaches, i.e., the use of remote technology to monitor the behavior, location, or physiology of individuals (Cooke et al. 2004). One of the most common forms of biotelemetry makes use of GPS or other transmitters to collect spatial location data of free-ranging animals. Although the efficacy and precision of this technology has improved greatly over time, the information obtained is fundamentally unchanged as a collection of time-stamped x-y coordinates. The umbrella of biotelemetry also includes various forms of what are collectively known as biologgers. Typically deployed in addition to spatial transmitters or recorders, these loggers can record several types of data either from the animal or from its immediate environment, e.g., velocity or ambient temperature (Box 1; Rutz and Hays 2009).

#### Box 1. A summary of how various biologgers function and what data they collect.

##### Accelerometers

(one, two, or three-axis) measure rates of acceleration along perpendicular axes using a piezoelectrical sensor. Somewhat analogous to a spring that compresses and expands when a force is applied, this sensor records voltages as velocity changes through time (Brown et al. 2012). These voltage changes are then converted to quantitative measurements of acceleration. These units can record at very high frequencies, often 1 – 100 Hz, providing extremely high-resolution data on the movement of an animal’s body. Typically, before being deployed on wild animals, preliminary truthing trials are conducted where the behavior of an accelerometer-equipped captive or otherwise directly observable individual is recorded continuously. These behavioral descriptions are then matched to the accelerometer’s output at the same time, and one of several machine learning algorithms can be trained to identify behavioral states in acceleration data obtained from wild individuals to obtain continuous time budgets for free-ranging wildlife (Leos-Barajas et al. 2017).

##### Acoustic

loggers are miniaturized audio recorders, typically attached to a collar or other mounting device. They can be programmed to various duty cycles, either running continuously, or set to record e.g. for 10 s every 10min to extend their temporal coverage and data storage abilities. The primary function is often to record vocalizations or other sounds of the animal wearing it, but they can also be used similarly to accelerometers to identify behavioral states based on non-vocal sounds characteristic of different movement gaits or behaviors (Stowell et al. 2017).

##### Activity

loggers record similar data to accelerometers but use a physical design rather than recording voltage changes. Typically involving a miniscule tube comprising a mobile internal component and sensors at either end, these loggers record an impact every time the mobile component touches one end of the tube; summing these impacts over a period of time gives an overall measure of activity during that period. If positioned on a collar (sometimes with a second perpendicular axis), these loggers can record vertical head movements, such as those of an ungulate while foraging, to derive head up – head down intervals.

##### Magnetometers

measure the direction and magnitude of magnetic field strengths. They are often used similarly to accelerometers to derive information about body position and movement. Given the planet’s magnetic field and the spatial location of the animal, the direction and movement of the animal can be derived from changes in the magnetic field sensed by the logger (Chakravarty et al. 2019).

##### Posture

sensors work similarly to activity loggers using a column with a mobile internal component, but rather than recording a number of events over each time period, they use gravity to continuously measure how the sensor is positioned in space (similar to a line level used in construction). They are often connected to spatial transmitters such that the telemetry signal changes depending on the position of the sensor. For species with characteristic resting and active posture (e.g., a bird of prey that roosts upright but flies with a horizontal posture; (Penteriani et al. 2011) this allows researchers to identify the activity state for each position recorded.

##### Pressure

loggers primarily serve to measure dive depth in aquatic and marine animals. With a known relationship between water depth and pressure, units of pressure (mmHg, kPa, or atm) can be converted to m below the surface, which is highly informative of the movement and foraging patterns of marine species like penguins, pinnipeds, whales, and diving seabirds (Bestley et al. 2013). They can also be used to estimate altitude of aerial species, particularly when collecting very fine-scale data or studying small-bodied species (Dreelin et al. 2018) as these loggers can be made substantially smaller than GPS tags. The GPS units deployed on birds or other flying species generally can record position in three dimensions, including altitude, so the two are rarely deployed together due to this redundancy.

##### Temperature

loggers (thermosensors, thermocouples) can be deployed on an animal or, in rarer occasions, internally within the body cavity to record body temperature. The latter are often combined with heart rate loggers to measure metabolic or physiological state over time (Menzies et al. 2020). External temperature loggers are useful for species with refugia such as nests or burrows, particularly those in cold environments, where a record of the temperature immediately around the animal can indicate resting vs. active periods or number of nest visits. In semi-aquatic species like beavers or waterfowl, time on land vs. time underwater can be distinguished via rapid changes in temperature as bodies of water are often not the same temperature as the air surrounding them. Thermosensors are relatively small and have low energy requirements compared to other biologgers, and often temperature loggers are by default included with other loggers like accelerometers (Studd et al. 2019) or depth sensors (Patterson et al. 2019).

##### Video

loggers are similar to acoustic loggers aside from their recording of visual data as opposed to acoustic. The location of the camera lens on the animal determines what sort of data can be collected and analysed: collar-mounted cameras typically show the lower jaw of the individual, useful for feeding events and identifying forage species (Lavelle et al. 2015). Backpack– or shoulder-mounted cameras facing over the top of the head can provide more of a first-person view of the animal’s daily activities (Naganuma et al. 2021). It is not possible to position these cameras in such a way as to record the entire body of the animal, so unlike a motion-activated camera or direct video recordings of an animal, behaviors are interpreted from the animal’s perspective.

While behavioral states and decisions can be estimated from movement data, these inferences are solely based on changes in the position of an animal over time and are generally less precise and at broader spatiotemporal scales. The second category permits more directly drawn inferences, e.g., three-dimensional accelerometers can identify specific behaviors to the precision of individual bite rates (Iwata et al. 2012). This precision (often multiple observations per second) leads to much higher battery and data storage demands, so these loggers generally cannot be deployed as long as GPS devices. Technological developments in the miniaturization of batteries and other technologies have made it possible to deploy transmitters of all kinds on smaller species and for longer periods of time (Kays et al. 2015).

### Foraging and habitat selection

If foraging is understood to include the seeking out and acquisition of resources, one must consider the availability and location of resources available to be foraged. Seeking and acquisition implies that animals make decisions about movement towards resources, and the choice to exploit or disregard resources that an individual comes across – analogous to the process of habitat selection. Such a framework is foundational to modern understandings of habitat selection; the third and fourth orders of selection identified by Johnson (1980) refer to the usage of specific habitat types or locations within the home range and the selection of individual food items within a feeding patch, respectively. While the latter would far more likely be described as foraging and not habitat selection, these processes exist along a spectrum and are not disparate concepts. As such, for the purposes of comparing direct observations and remote approaches, which are more likely to use the lens and language of habitat selection, we broadened our definition of foraging to be more inclusive.

### Objectives

Remote methods could provide valuable and complementary insights to behavioral ecology by expanding the scope of research to temporal and spatial scales that are not feasible through direct observation. In an effort to synthesize across disciplines, taxa, and methods, we performed a literature review of foraging behavior studies. We had two primary objectives:

1. Compare direct observational and remotely sensed approaches to studying foraging behavior across spatial and temporal scales.
2. Highlight what insights each approach offers and evaluate their underlying assumptions.

## Methods

### Literature search and refining

We used Web of Science to find research on the foraging behavior of wildlife. Foraging behavior is an expansive research field, so we narrowed our search based on several criteria to compare direct observations and remote approaches (summarized in Table 1). Our focus was how various approaches to studying foraging differ in their spatiotemporal scale and the information they can provide about wildlife behavior. We omitted studies of captive animals and provisioned populations, but this still retained an unmanageably large number of papers. In the interest of comparing direct observations to remote approaches in the same systems, we also excluded studies of marine species (which are rarely directly observed) and invertebrates (most of which are too small to deploy remote technologies). After these restrictions, 2912 studies of wild terrestrial vertebrate foraging behavior remained. We further limited the search to publications from the year 2010 onward as many of the remote approaches we were interested in were relatively recently developed. The series of restrictions then returned 729 studies, 125 of which we later excluded as non-behavioral research, e.g., measured hormone concentrations. The final statistical analysis thus included 604 studies (Figure 1).

**Figure 1.**
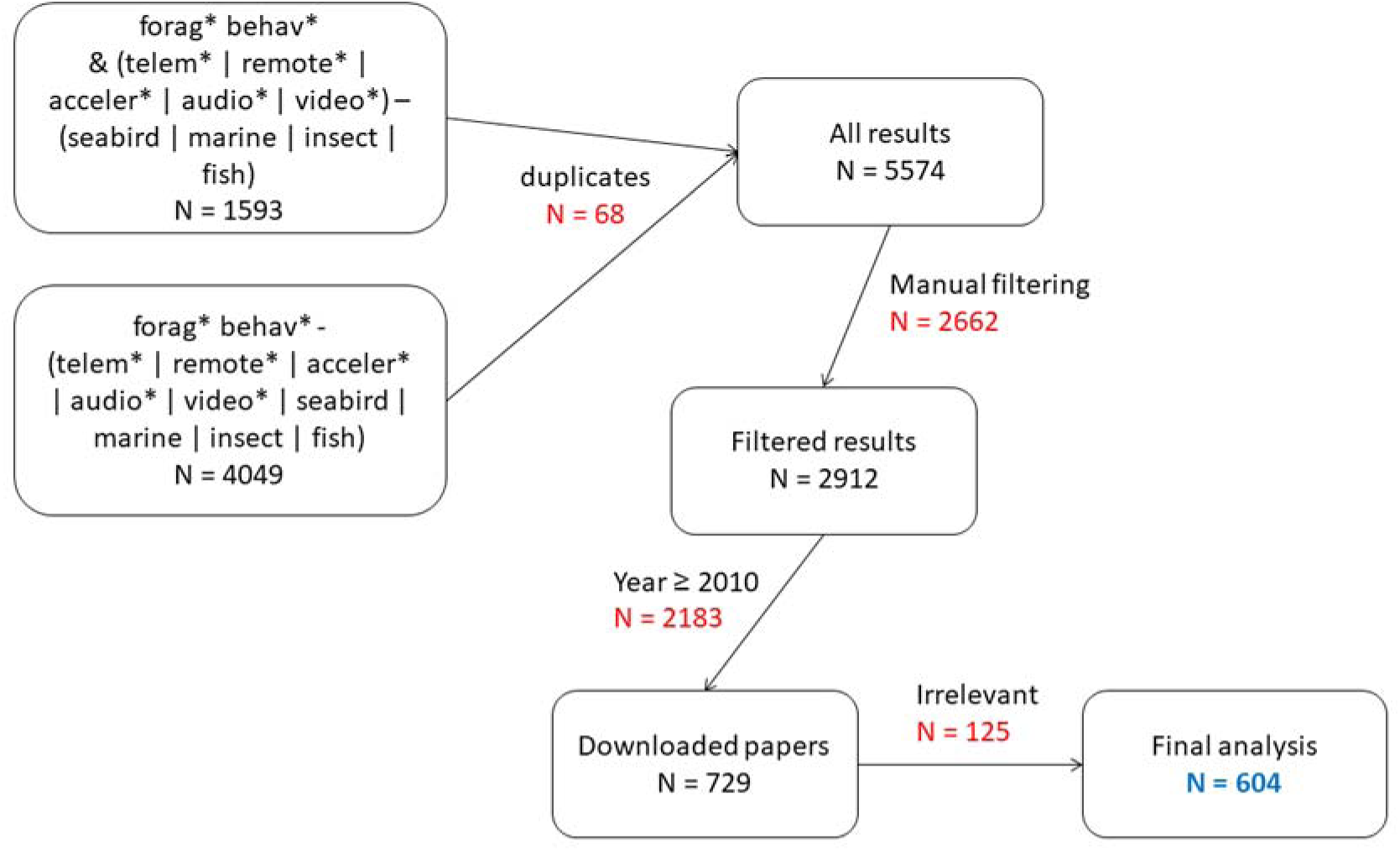
Overview of the literature search (conducted 12 October 2021) and selection process to obtain the final dataset of 604 studies of foraging behavior analysed in this review.

**Table 1.**
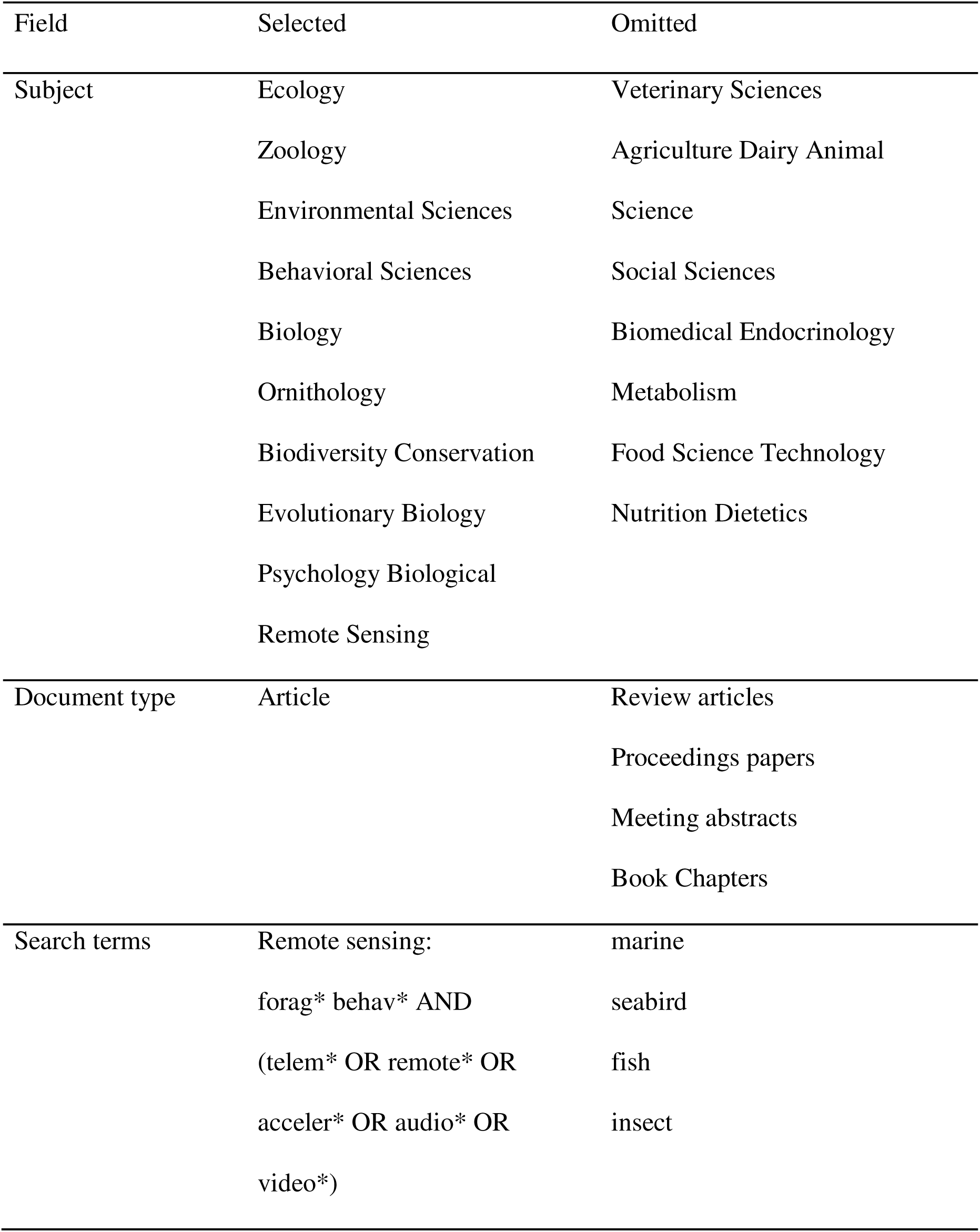
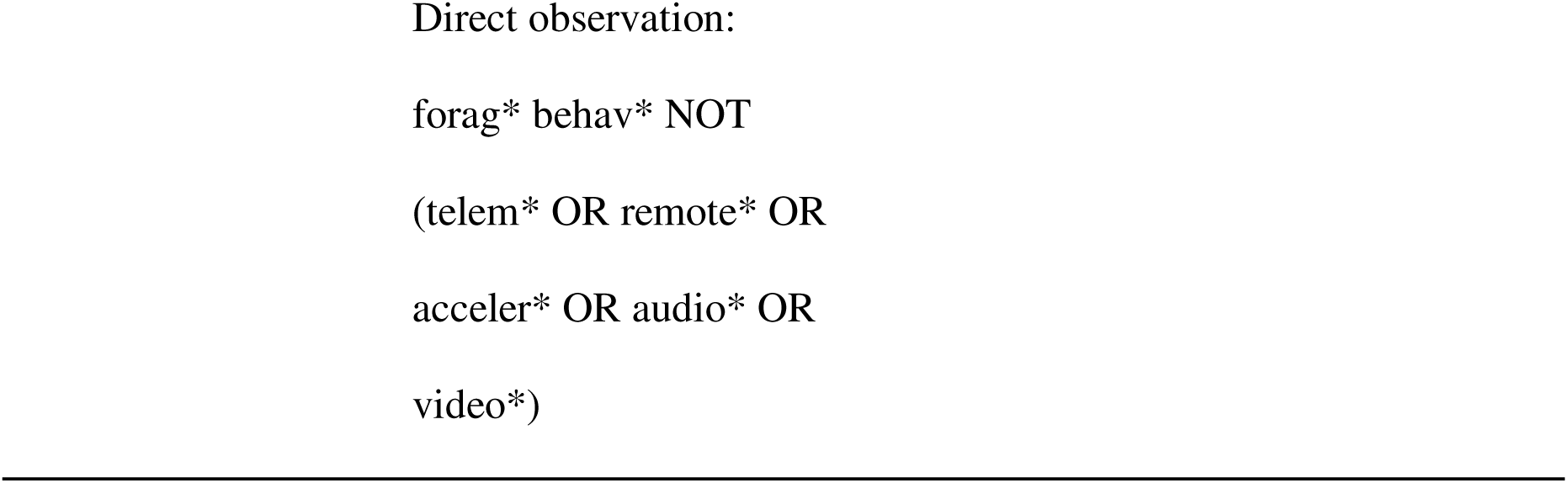
Literature review criteria used in Web of Science on 12 October 2021. Search terms were given for all fields, i.e., anywhere in the title, abstract, or keywords of the study.

We used two separate literatures searches to collect studies of foraging behavior using direct observation separately from studies using remote approaches (Table 1). Each paper was thus pre-categorized as either observational or remote based on the search terms through which we identified it. However, through the process of reviewing papers, it became clear that this dichotomy did not adequately characterize the breadth of methods and approaches used by researchers. We recorded the research method as stated by the authors, e.g. radio telemetry, GPS, direct observation, camera trap, listing multiple if the study used more than one approach. After reviewing all papers, we used a k-means cluster analysis on the 35 unique methods to determine five meaningful categories of research methods (details in Appendix 2) rather than our initial dichotomy.

### Data extraction

For each of the n = 604 studies, we primarily extracted methods data and basic metadata including the study species and Order. We did not record effect sizes or other results, as our interest was not in conducting a meta-analysis, but rather in ascertaining how various approaches studied foraging across scales. We considered four metrics in this review: spatial extent, temporal extent, temporal coverage, and behavioral stage.

Scale comprises an extent (the entirety or size) and a grain (the unit measurement of discrete units (Wiens 1989). We extracted the temporal and spatial extent of each study, such as a one-year long study (temporal extent) where the animals ranged over 10 km^2^ (spatial extent). We also recorded the amount of time throughout that extent during which data were collected as a proportion of the total (temporal coverage). Recording observation period allowed us to differentiate between studies with the same total duration: researchers could spend 8 hours every day for a month making focal observations, or only observe animals for one hour each week during that same month. The temporal extent would be identical for each, but the temporal coverage or actual observation period differs by an order of magnitude.

We were interested not only in the spatial and temporal scale of foraging studies, but also in the type of information and the insight offered by each, characterized here as behavioral stage. Our working definition of foraging, “the seeking out, acquisition, and consumption of food resources,” comprises three stages of progressively finer scales. Following Johnson (1980), we defined three stages: habitat selection, searching, and feeding. We recorded which of these three stages, either alone or in combination, each study investigated. By comparing which stages were more commonly studied by different methods of data collection, we can determine not only if direct or remote approaches differ in their spatiotemporal scale, but also whether they offer differing ecological insights.

### Inference of missing values

Some papers did not report one of our metrics of interest. Spatial extent was not stated in ∼38% of papers (n = 230), and this was more common for direct observations (n = 158, 57%) than for other research approaches. Some of these studies (n = 38) reported multiple sites (e.g., “behavioral trials were conducted at 8 locations”, “camera traps were deployed at six points”). In the interest of not excluding nearly two-fifths of our data, we imputed a value of 100 m^2^ for studies with missing spatial extent, representing a 10m x 10m square as an approximation of the field of view of a researcher or camera. For studies conducted at a number of sites, we multiplied this number by 100m^2^ to get the total extent. Temporal extent was absent from only 3.6% of studies (n = 22), so we simply omitted these studies from our temporal analyses.

Temporal coverage also required some inference depending on research approach. Nearly half of the studies we found (n = 272, 45%) reported observations not as a continuous measure in minutes or hours, but as point observations at intervals, e.g., GPS collars recording locations every two hours or instantaneous scans of behavioral state every 30 minutes. Remote studies were more likely to report discrete time points (85%) than were direct observations (22%). We elected to treat each instantaneous estimate as 1 second and converted these values to an observation period.

### Statistical analyses

We performed all statistical tests in R v4.1.0 (R Core Team 2021). For our analyses of spatial and temporal scale, we modelled spatial extent, temporal extent, and temporal coverage as a function of research approach, which was a categorical predictor with five levels representing the five categories determined through our cluster analysis. Spatial and temporal extent were both non-normally distributed and so were log-transformed prior to analysis; temporal coverage was left untransformed as a proportion.

For our analysis of behavioral stages, we took a more descriptive approach rather than building a predictive model. We calculated the proportion of studies within each category that investigated habitat selection, searching, and feeding behaviors. We used these proportions to assess whether certain approaches were better suited to, or more likely to be used for, studying different behavioral stages within the overall field of foraging behavior.

## Results

### Categories of approaches

The various methods of studying foraging behavior identified in this review (n = 35) were classified via *k*-means clustering (Appendix 2) into five categories of approaches: direct observation, manual tracking (via snow, footprints, etc.), biologgers, remote audio-visual, and remote spatial (Table 2). According to our initial framework of an observational vs. remote dichotomy, direct observations and manual tracking would be collapsed into a group opposite biologgers, remote audio-visual, and remote spatial approaches. Such a proposed dichotomy fails to adequately describe the variation in these approaches: the differences within these two groups are as large, if not larger, than the differences between the two. We thus dropped the binary categorization of approaches and adopted these five categories in all analyses. Our review included 324 direct observation, 214 remote spatial, 105 manual tracking, 68 biologger, and 54 remote audio-visual studies of foraging behavior. Some studies incorporated multiple approaches, so these values sum to more than the *n* = 604 total studies we reviewed.

**Table 2.**
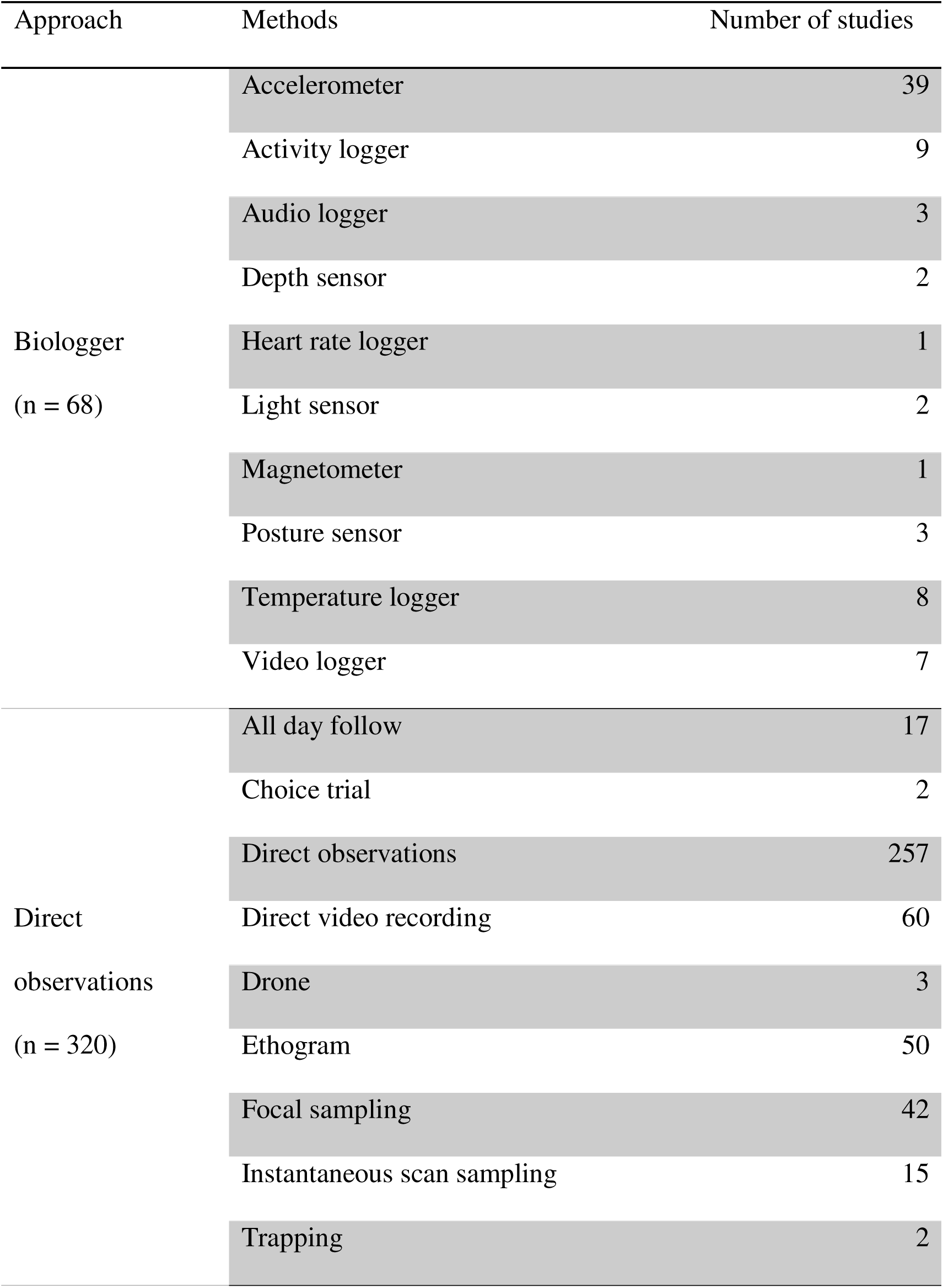
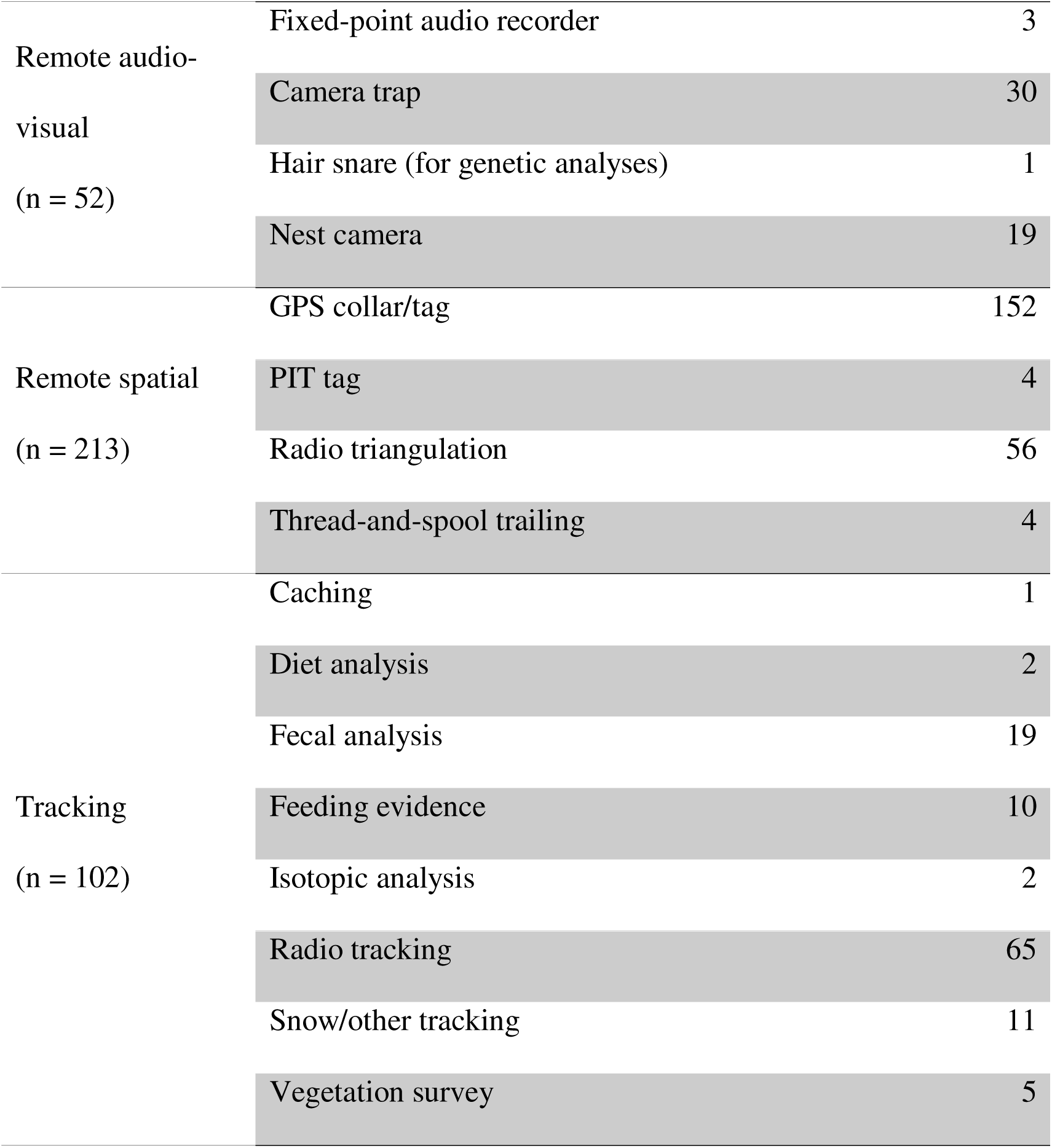
The 35 distinct methods identified in this literature review of n = 604 studies of foraging behavior were sorted into five approaches. The number of studies included in each approach sum to more than 604, as some studies used multiple approaches in their research.

### Spatial and temporal scale of approaches

Studies with larger spatial extents tended to have larger temporal extents across the full dataset (Figure 2), but this relationship varied somewhat when looking at each of the five approaches individually (Table 3). Approaches tended to occupy distinct spatial and temporal ranges from one another (Figure 3). Direct observations and remote audio-visual studies had the smallest spatial extents, followed by biologger and tracking, with remote spatial studies covering the largest area, as might be expected (Figure 4).

**Figure 2.**
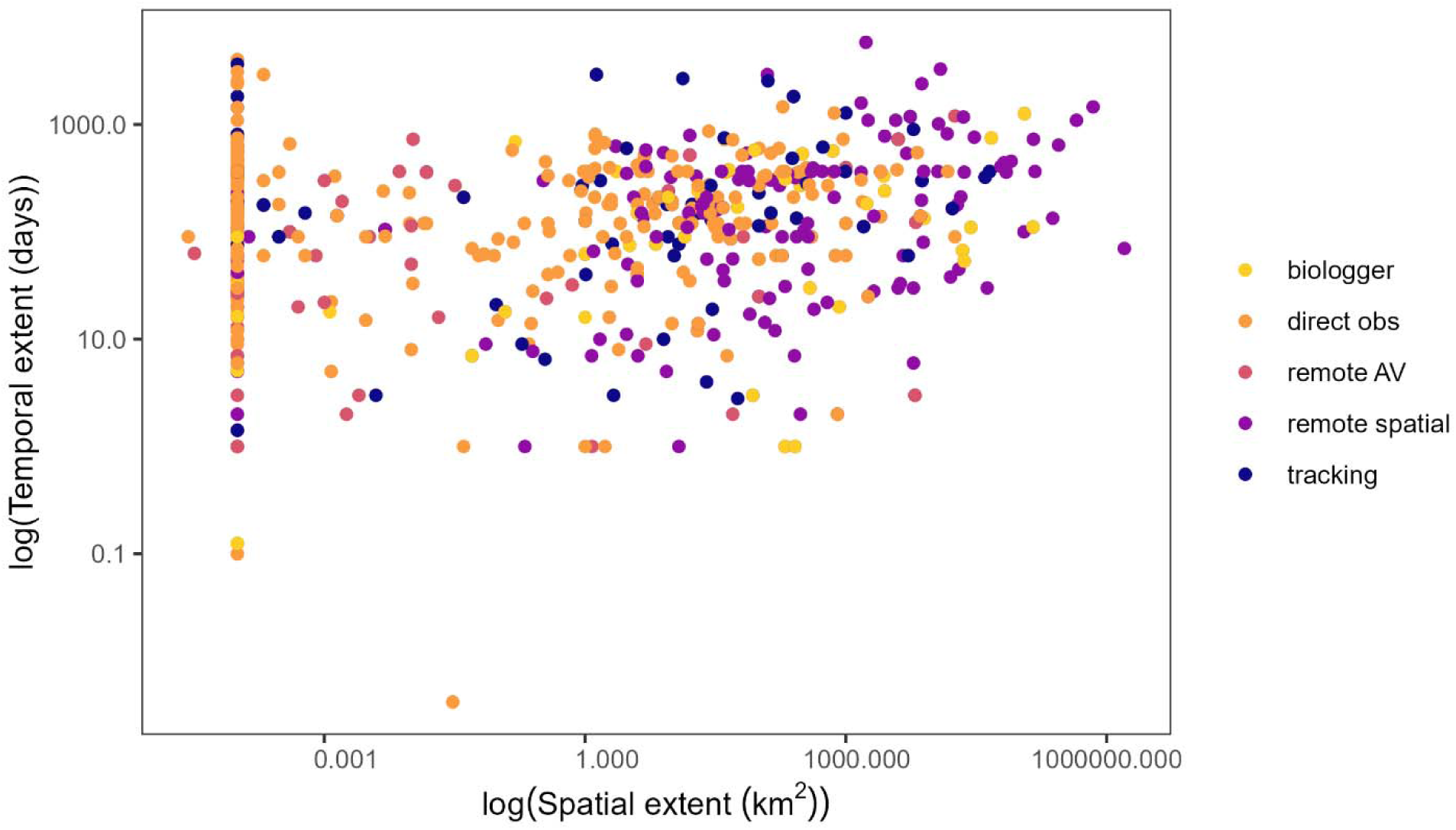
Scatter plot of spatial and temporal extent of 604 studies of foraging behavior; both metrics of scale were log_10_ transformed for visualization purposes and in statistical analyses. Colors indicate one of five categories of approaches for methodology of each study (biologger *n* = 67, direct observations *n* = 312, remote audio-visual *n* = 53, remote spatial *n* = 210, tracking *n* = 98).

**Figure 3.**
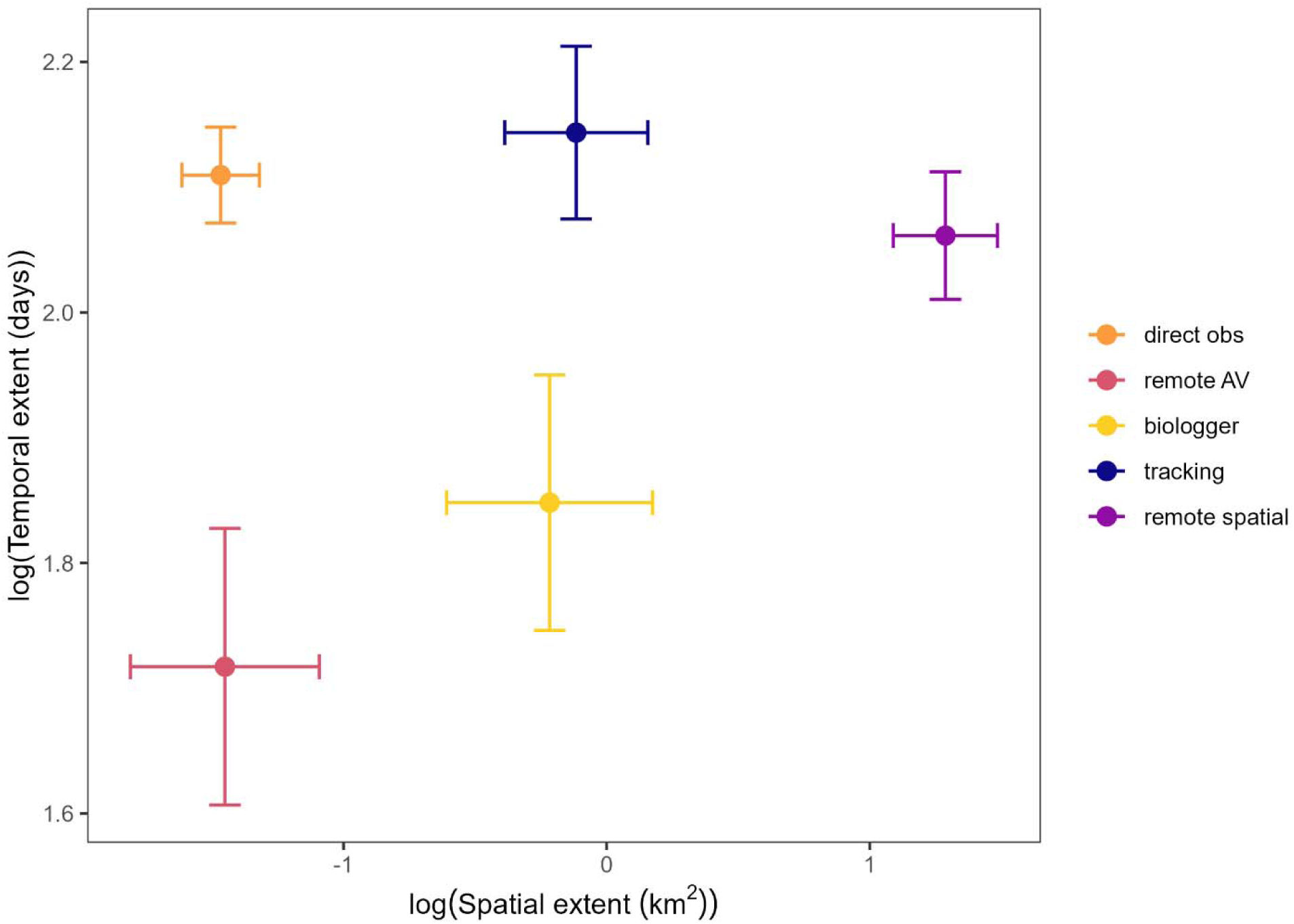
The average ± standard error for the spatial and temporal extent of studies falling within five categories of approaches to studying foraging behavior (biologger *n* = 67, direct observations *n* = 312, remote audio-visual *n* = 53, remote spatial *n* = 210, tracking *n* = 98).

**Figure 4.**
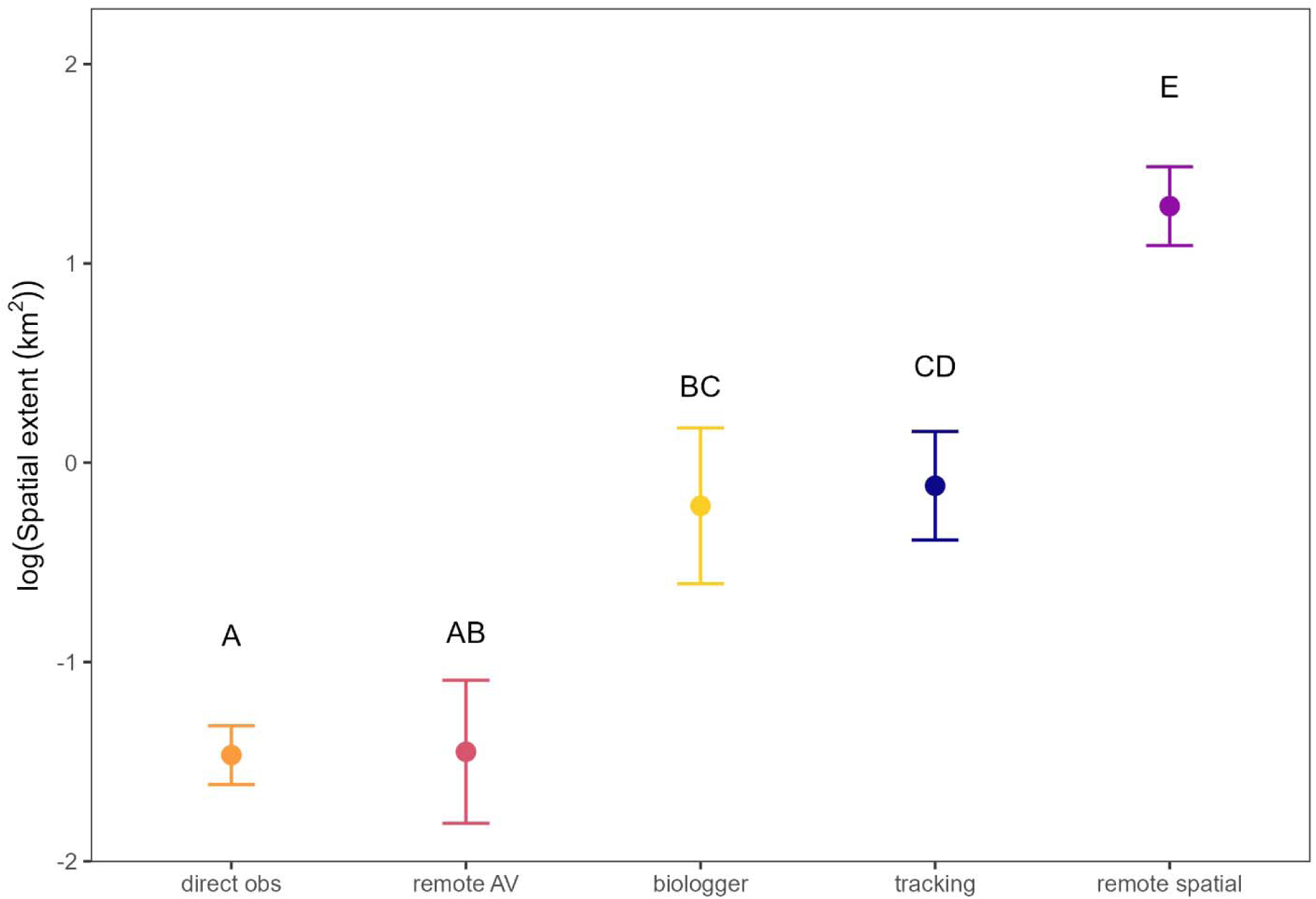
Average ± SE spatial extent of studies of foraging behavior, separated by methodological approach (biologger *n* = 68, direct observations *n* = 324, remote audio-visual *n* = 54, remote spatial *n* = 214, tracking *n* = 105). Letters indicate significant differences between approaches (α = 0.05).

**Table 3.** Regressions of log_10_(temporal extent) as a function of log_10_(spatial extent) for all studies, and within each category of approaches to determine whether the relationship varied by study methodology. These regressions are not intended to imply a causal relationship between the two; we arbitrarily selected temporal extent to be the dependent variable in these models.

| Approach | *n* | Estimate ± SE | *t* | *df* | *p* | *r* |
| --- | --- | --- | --- | --- | --- | --- |
| All | 740 | 0.035 ± 0.009 | 3.95 | 738 | < 0.001 | 0.14 |
| Biologger | 67 | 0.082 ± 0.031 | 2.68 | 65 | < 0.001 | 0.32 |
| Direct observation | 312 | 0.007 ± 0.015 | 0.50 | 310 | 0.62 | 0.03 |
| Remote audio-visual | 53 | 0.075 ± 0.042 | 1.79 | 51 | 0.08 | 0.24 |
| Remote spatial | 210 | 0.071 ± 0.017 | 4.11 | 208 | < 0.001 | 0.27 |
| Tracking | 98 | 0.015 ± 0.026 | 0.57 | 96 | 0.57 | 0.06 |

Remote audio-visual studies had the shortest study durations or temporal extents. Biologger studies had an intermediate length, while the remaining approaches had similarly long total study durations (Figure 5). Temporal coverage, or the proportion of the temporal extent during which data were collected, had a generally opposite trend to temporal extent: approaches with larger temporal extent tended to have lower temporal coverage. Remote spatial and direct observation studies both had the lowest coverage, while biologgers and remote audio-visual studies had much higher coverage, up to and including 100% (Figure 6).

**Figure 5.**
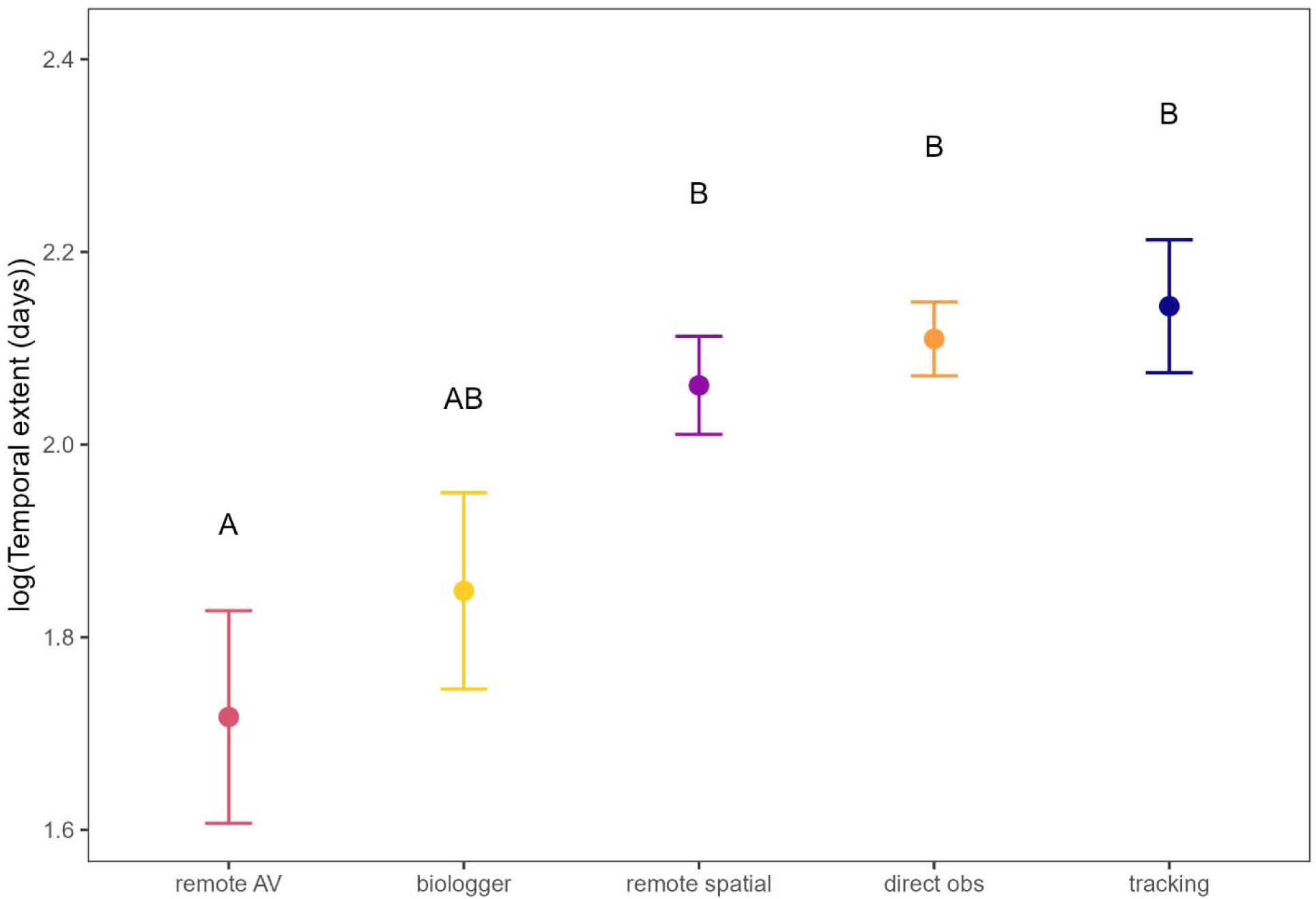
Average ± SE temporal extent of studies of foraging behavior (log-transformed number of days) separated by methodological approach (biologger *n* = 67, direct observations *n* = 312, remote audio-visual *n* = 53, remote spatial *n* = 210, tracking *n* = 98). Letters indicate significant differences between approaches (α = 0.05).

**Figure 6.**
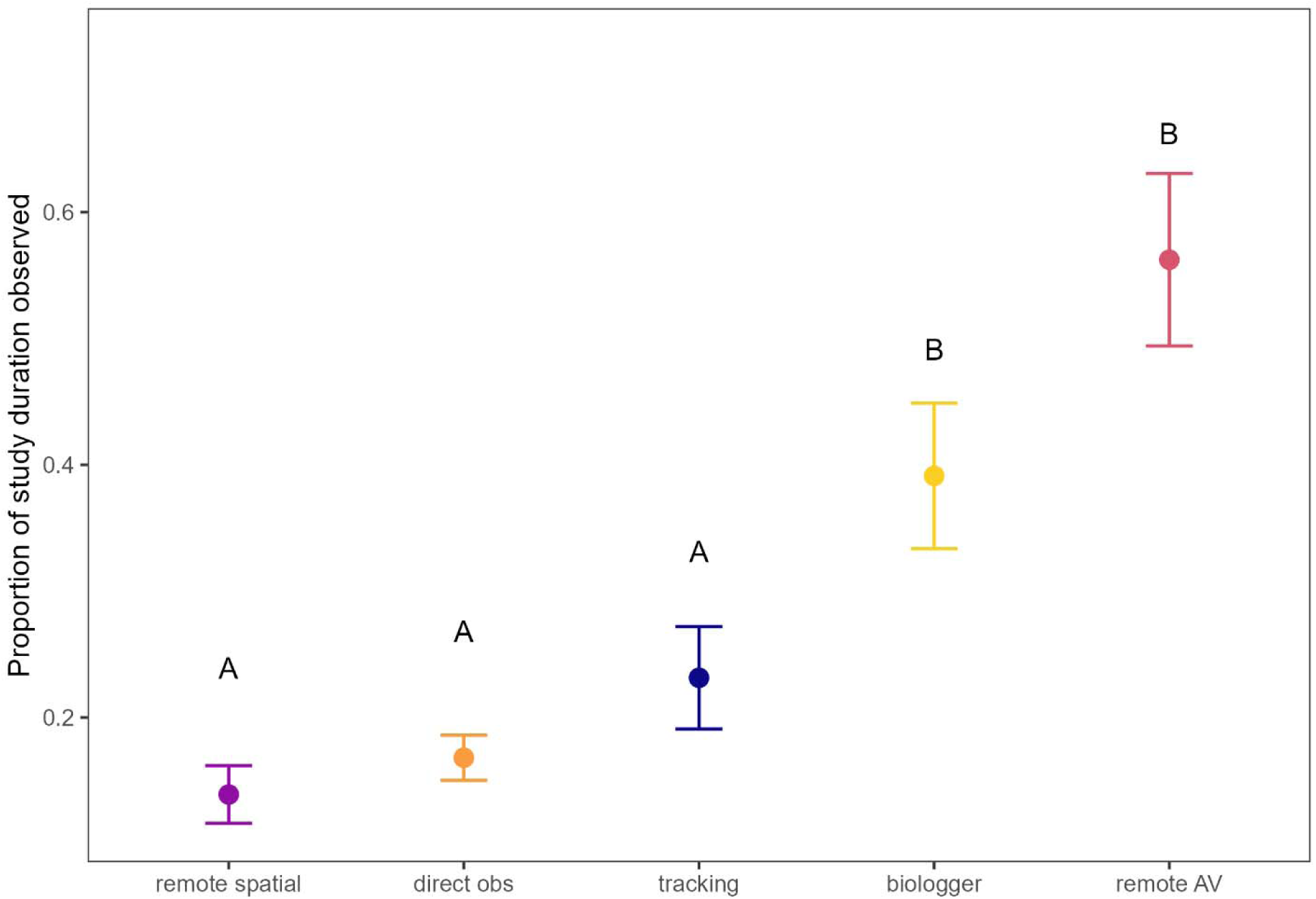
Temporal coverage, or the proportion of the total study duration (temporal extent) that was spent observing the study species (biologger *n* = 66, direct observations *n* = 279, remote audio-visual *n* = 41, remote spatial *n* = 198, tracking *n* = 86). Letters indicate significant differences between approaches (α = 0.05).

### Use of observational and remote approaches at each behavioral stage

Most studies in this review looked at more than one stage of the foraging process. As such, when summarising the proportion of papers investigating each stage, the sum of the percentages exceeds 100% for all approaches (Figure 7). Direct observational studies were concerned primarily with finer scale feeding (85%) and searching (90%) behavior, whereas remote spatial approaches were dominated by habitat selection (94%). The most common stage investigated by biologger and remote audio-visual studies was searching (90% and 70% respectively), while tracking studies were relatively evenly distributed across all behavioral stages (59% feeding, 68% searching, 58% habitat selection).

**Figure 7.**
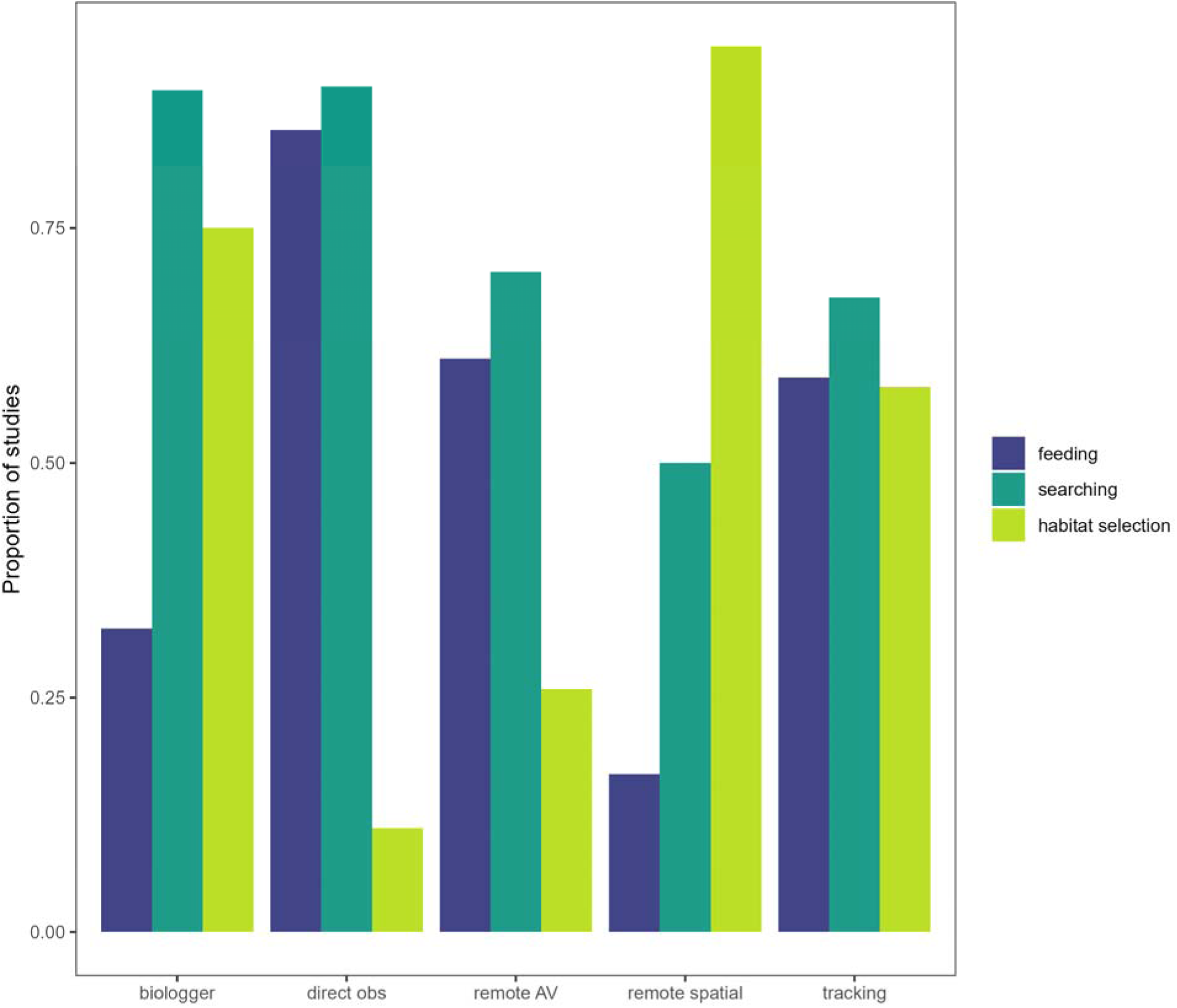
The proportion of studies within each approach to studying foraging behavior (biologger *n* = 68, direct observations *n* = 324, remote audio-visual *n* = 54, remote spatial *n* = 214, tracking *n* = 105) that investigated each of three sequential and nested stages in the foraging process: active feeding behavior, searching for food, which precedes this feeding, and habitat selection that precedes searching behavior. Most studies looked at more than one stage of the process, hence the proportions within each approach add to more than 100%.

## Discussion

### Remote vs. direct observations: a false dichotomy

At times disciplines in biology operate in their own silos. Even within disciplines the different practical methods used to test hypotheses can contribute to further segregation. We perceived such separation might exist between behaviorists that typically make direct observations of their study animals and those that monitored the animals remotely. Our premise, however, was oversimplified. We intended to spotlight our premise using foraging as a case study comparing direct and remote observations, but our literature review illustrated that our premise was flawed, and failed to identify the multitude of approaches to studying foraging behavior. We replaced our initial dichotomous categorization of direct vs. remote observations with five groups of approaches, defined not by technology, but rather by the type of information provided and the underlying assumptions of each approach. Under our revised framework, direct observations comprised any study where animals were watched live by researchers, whether data were recorded in the moment, or if animals were video recorded and these videos later scored (e.g. Austin and Ramp 2019). The confidence in behavioral assignment and species identity, and the type of information provided, is identical in these cases regardless of whether observations were mediated by a camera, binoculars, or any other optical aid. Forcing studies into a binary classification based on the use of technology, rather than on the data produced and the inferences permitted by those data, is not an effective way of analysing the published literature and elides some more meaningful distinctions between studies of foraging behavior.

### Scale – extent vs. coverage

Spatial and temporal scale of studies of foraging behavior varied dramatically across the studies (n = 604), from a singular point to more than 1.6 million km^2^ (Wallau et al. 2010; Eisaguirre et al. 2019), and several minutes to more than a decade (Laundré 2010; Walsh et al. 2013). The five approaches differed in their typical range of spatial and temporal extent, although there was some overlap. Remote spatial studies covered the largest spatial extents, while direct observational and remote audio-visual approaches had the smallest areas. These latter approaches were also the most likely to fail to report an area, and thus included more of the 100m^2^ interpolated extents. We posit that this is reflective of a methodological reality, rather than an artificial consequence of our analysis; if authors do not report the area over which the study took place, spatial extent was presumably immaterial to the study and thus unlikely to encompass any significant area.

While spatial extent and temporal extent were weakly correlated across studies (log-transformed r = 0.14, t= 3.974, df = 738, p < 0.001), studies covering smaller spatial areas did not necessarily have shorter temporal durations, although remote audio-visual studies tended to be low for both metrics. This relationship varied across the five approaches. Remote spatial approaches had the largest spatial extents, but intermediate temporal extent; tracking studies had the longest duration (Figure 5), but intermediate area (Figure 4).

Studies with large temporal extents, i.e., those with very long study durations, did not necessarily have more observations of the study subjects through time. Temporal coverage, which reflects the proportion of the total study duration during which data were actively being collected, was inversely related to temporal extent (Figure 6). The negative association occurred also when looking at temporal observation periods in isolation, and as such is not solely a consequence of increasing denominator in the temporal coverage ratio. It is somewhat inevitable that direct observational studies would have quite restricted temporal coverage throughout extensive study durations, given human limitations. Even established ethological programs where researchers continuously observed study subjects during daylight, 27.5 days per month, had an effective coverage of only 59% across a 20-month study duration (Lucchesi et al. 2020). While the data obtained from work such as this are extremely valuable, this represents an enormous investment of human time and labour over a multi-year period that would be an insurmountable barrier for most researchers.

Temporal extent is likely more reflective of practical or logistical constraints of a given study. Direct observations and remote audio-visual studies have similar spatial extents, but observational studies tend to occur over much longer timescales compared to research using camera traps which are relatively restricted in time. Parallel to these, biologger and tracking studies both range over comparable spatial areas, but biologgers have shorter temporal extents, likely due to technological limitations. Tracking may either involve no animal-borne technology, or simply a radio transmitter; biologgers such as accelerometers or depth sensors have higher energetic and data storage requirements than spatial transmitters and so their deployment time is often much shorter than that of GPS or VHF tags (Krause et al. 2013), although this discrepancy is improving over time (Korpela et al. 2020).

Temporal extent does not necessarily indicate the amount of observations of a given study, which is when temporal coverage may be a more valuable metric to consider the ‘efficiency’ of an approach. Temporal coverage for observational and tracking studies is relatively simple to calculate: the sum of hours or days spent following the study subject, as a fraction of the total number of days between the first and last observation. Temporal coverage is more complicated for studies that collect data not continuously, but as repeated point estimates. We calculated temporal coverage for GPS collars collecting fixes e.g., every hour using a 1-second estimate for each fix. As the interval between data points shortened, however, observations became more continuous. Most biologger studies had devices set to sub-second intervals, up to 50 or 100 Hz (Byrnes et al. 2011; Chakravarty et al. 2019), which we treated as continuously collected data. Some studies programmed loggers to record very fine scale temporal data only at subsamples through the deployment period to extend their lifespan, e.g., 16s of 3.3 Hz sampling once every 10 min (Spiegel et al. 2015), which works out to a temporal coverage of 2.67% (16 s / 10 min).

Observation periods for remote audio-visual studies are also somewhat subjective. It could be argued that these devices have 100% temporal coverage, as they are continuously collecting observations during the entire deployment time given the camera/audio recorder is turned on. In contrast, a case could be made to include only the time when a focal subject is being recorded as observations, excluding videos, photos, or audio files where no subjects are present. We elected to follow the latter criteria in this review, treating each photo observation from motion-activated still photography (Srbek-Araujo et al. 2017) as a 1-second observation similar to GPS fixes (above).

While 1 second per GPS fix is a reasonable facsimile for studies where points were analysed as discrete observations, some movement models draw inferences for the intervals between recorded point locations, e.g. Hidden Markov Models (McKellar et al. 2015), integrated step-selection analyses (Wilson et al. 2021) rather than analysing each fix as an instantaneous observation (Beumer et al. 2020). A claim could be made for considering this 100% temporal coverage, but interpolating of movement and activity between fixes is more analogous to periods of non-visibility in observational studies, and as such we did not make this assumption in our review. Much like a clear, non-subjective delineation cannot be drawn between “remote” and “direct” methods of observation, it is similarly challenging to objectively define the period during which individuals are being observed in many studies.

### Behavioral stages of the foraging process

The inferences provided by each approach to studying foraging behavior tended to align with our three behavioral stages of the foraging process (habitat selection, searching, and feeding). Direct observations and remote audio-visual studies were primarily concerned with feeding and fine-scale searching behavior, as the restricted spatial scale of these approaches made larger scale movements and searching behavior difficult to follow. Tracking studies were the most evenly distributed across stages, likely due to the diversity of methods in this approach: some are better suited for measures of feeding (e.g., fecal analysis), while other methods within this approach better suit searching and habitat selection analyses (e.g., tracking, thread trailing).

The majority of biologger studies focused on the searching stage, as searching behavior is easier to assign from biologger output. Data from accelerometers and magnetometers are typically divided into behaviors such as resting, foraging, and higher activity behavioral states. The minute bodily motions associated with feeding behavior are generally more difficult to identify from these data than the more distinctive signature of searching and foraging (but see (Studd et al. 2019). Biologger studies also included measures of habitat selection, but this is primarily because biologgers tended to be deployed simultaneously with GPS or other remote spatial technologies that allowed for this broader-scale analysis. Similar to biologgers, remote spatial approaches can infer behavioral states from movement patterns and identify foraging behavioral states via area-restricted search, path tortuosity, or step length and turning angle models (Beumer et al. 2020). Behavioral states, however, are assigned as an overall period of time rather than specific behaviors. For example, an animal could be assigned a “foraging” state between two consecutive hourly GPS fixes, but this does not mean the animal was continuously foraging for that entire hour. While remote spatial studies did tend to investigate searching behavior (50%) more than feeding (16%), habitat selection was by far the most common stage of the foraging process studied by remote spatial approaches (94%, n = 202 of 214 studies).

We considered habitat selection as part of foraging in this review primarily to ensure remote approaches were meaningfully considered. After our analysis, we maintain that foraging and habitat selection are integrated phenomena along a spectrum of behavior. Both comprise decisions about movement and resource selection. We note this as an example of seemingly disparate behavioral subfields that may be isolated more due to human constructs rather than intrinsic ecological differences.

### Conclusions: assumptions and caveats

While remote inferences require assumptions to be made about the relationship between data output and the actual behavior, classical behavioral research is not devoid of assumptions. Assumptions are integral to all ecological research but are often left unstated or implicit and taken for granted, particularly for well established research protocols that are ingrained in the field. To observe individuals in the wild, there must be sufficient light and relatively open habitat to see animals behave. If we rely solely on direct observation, we are unable to know whether behavior differs at night, in dense habitats, or in inaccessible marine (Johnson et al. 2009) or subterranean contexts (Martín et al. 2021), among others. Observer effects are a more frequently noted potential influence on wildlife behavior. Habituation protocols can alleviate some of these concerns (Johns 1996), particularly when the success of habituation is validated through cross-sectional or longitudinal comparisons (Gazagne et al. 2020). Nonetheless, observer effects can persist in subtle ways, as individual animals can differ as to the rate of and capacity for habituation (Allan et al. 2020), and even ostensibly habituated populations have been shown to behave differently when human observers are absent (Titus et al. 2015). Outside of its application as a research tool, habituation to humans can occur non-deliberately and aggravate human-wildlife conflicts (Shimozuru et al. 2020; Cui et al. 2021), though whether deliberate research– oriented habituation generalizes to other human contexts remains an open question (Powell et al. 2022). We raise these issues not to invalidate or deny the usefulness of classical behavioral observations: these studies have provided and continue to provide invaluable data on innumerable populations. But too often direct observations are held up as a paragon against which remotely-inferred behavior is compared and found lacking. No observations of nature are entirely complete, regardless of approach; it is only the scope and nature of their assumptions and insights that vary.

Different underlying assumptions in research methods can in fact lead to complementary insights. Studies using multiple approaches simultaneously are able to draw broader conclusions than those only investigating one aspect of foraging behavior, as the assumptions of one approach can be verified by another. In this review, 23% of studies used more than one of the five types of approaches identified here, and many of their conclusions hinged upon this multi-pronged strategy. Studies making use of multiple approaches can reconstruct a more complete representation of foraging behavior than those coming from only one perspective. For example, when investigating the foraging behavior of wetland-specialist waterbuck (*Kobus ellipsiprymnus*) during an expansion into historically avoided savannah habitat, fecal analysis indicated that digestibility and protein availability was lower in the savanna, yet GPS-accelerometer collars found that waterbuck spent less time foraging in savannah despite the poorer nutrition available (Becker et al. 2021). Demographic differences in which individuals made use of this novel habitat were also identified via motion-activated trail cameras. None of these approaches in isolation would provide the same overall narrative as considering them in concert. Similarly, the impact of non-native vegetation on the diet and foraging of Galapagos tortoises (*Chelonoidis porteri*) was estimated through GPS locations, ethological observations, and manual tracking (Blake et al. 2015). Fecal analyses tied to GPS tracks found increasing prevalence of invasive species in tortoise diet at higher elevations, but direct observations of feeding behavior saw no change in the proportion of invasive to native species consumed. Invasive plants were more abundant at higher elevations when vegetative communities were surveyed, such that the observed change in diet was not due to tortoise selection but rather forage availability. Any of these findings alone would not lead to the same overall conclusion as that obtained by considering them together, again demonstrating the value of applying multiple approaches to the same question. Ultimately, we posit that remotely-inferred behavioral approaches should not be seen as inferior to direct observation, but rather appreciated for their complementarity to classical approaches. Simultaneous application of new technologies along with historical practices can improve both the scope and granularity of our insights into foraging behavior in its entirety.

## Author contributions

*Jack G Hendrix*: Data curation, Formal analysis, Investigation, Methodology, Visualization, Writing – original draft, Writing – review & editing. *Eric Vander Wal*: Conceptualization, Project administration, Supervision, Writing – review & editing.

## Data availability

A list of all the papers collected as part of this review, as well as our final dataset and the code used to replicate analyses, can be found at https://github.com/jghendrix/foraging-tech/.

## Declaration of interest

The authors declare no conflicts of interest.

## Acknowledgements

This project was designed, conducted, and written in Ktaqmkuk, what is now known as the island of Newfoundland, on the ancestral homelands of the Mi’kmaq and Beothuk peoples. We are deeply grateful to several lab members whose suggestions and edits greatly improved the quality of this manuscript. We would also like to thank C. Walsh, P-P. Bitton, and G. Street who contributed to the topic of JGH’s comprehensive exam that was the inspiration for this project.

## Appendix

## Appendix 1: Defining foraging

In the interest of objectivity and considering differences between disciplines, we assessed how foraging was defined by researchers rather than coming up with a definition that ascribed to our own preconceptions. We searched Web of Science for the top 100 most-cited papers of foraging behavior and recorded any definitions of foraging that we could find. The most highly cited papers in foraging behavior all made use of direct observations, but given our research objective was to compare direct observations and remote approaches, we also sought out definitions from studies of remotely-inferred foraging behavior. We thus analysed the 50 most cited direct observation foraging papers as well as the 50 most cited remotely sensed foraging papers (Table A1; n = 89 total; 4 occurred in both, and 7 were not actually studies of foraging behavior).

More than one third of these papers (n = 33/89; 37%) included no definition of foraging at all. In fourteen papers (16%), foraging had a purely spatial definition: certain habitats or locations were referred to as foraging areas, and any visits or time spent in this habitat type was assumed to be foraging, without any actual observations of behavior. Thirteen papers (15%) used a movement-based definition of foraging, of which all but one used remote inferences. Turning angle, step length, area-restricted search, and the like were all used to assign behavioral states to movement data. Foraging was typically the label applied to intermediate movement rates, relative to either resting or other more directed movements. Eleven papers (12%) defined foraging as an event, typically a trip away from and returning to a central nesting location, e.g., for bees and seabirds, any time spent away from the colony was considered a foraging trip but the actual behaviors during this period were not explicitly studied. In five of the papers (6%), foraging was presented as a sort of default behavioral state within a fixed time budget: e.g., in a study of vigilance, any time not spent being vigilant was assumed to be spent foraging. Another five papers (6%) used the terms foraging and feeding synonymously, with no distinction between the two; three papers (3%) used evidence of feeding (proportion of seeds/fruit left from the initial total) as indicative of foraging. Only nine of the most highly cited papers on foraging (10%) provided explicit behavioral definitions of what constitutes foraging in their system.

Synthesizing the nine studies with actual behavioral definitions, we see foraging as a series of progressive stages involving the seeking out, acquisition, and consumption of food resources. These stages can be disconnected from one another in time: autumnal foraging by food caching species, where resources are collected and stored but not consumed until months later, still represents a form of foraging behavior.

Difficulties in definitions arise when setting the boundaries of what is included or excluded. Under our definition only food resources can be foraged. Desert animals seeking out water, or breeding birds searching for and collecting nesting materials, are not considered to be foraging for the purposes of our review. The boundary is also unclear in cases of parental provisioning. In such cases as songbirds collecting arthropods to feed their nestlings, the parents are not themselves eating these resources and benefitting from this energy. The parents could be considered foragers based on search and acquisition, or alternatively the nestlings consuming this provisioned food could be considered foragers themselves. For this review we favoured where possible the earlier stages of seeking out and acquisition as central to foraging; as such, nest-provisioning adults can exhibit foraging behavior (n = 43 studies) but sedentary nestlings cannot. In contrast, for the parallel life history stage of lactation in mammals, precocial offspring have agency in nursing, making decisions about when to nurse, or even from which adult (Packer et al. 1992) and are thus considered foragers in this review (n = 2 studies).

**Table A1.**
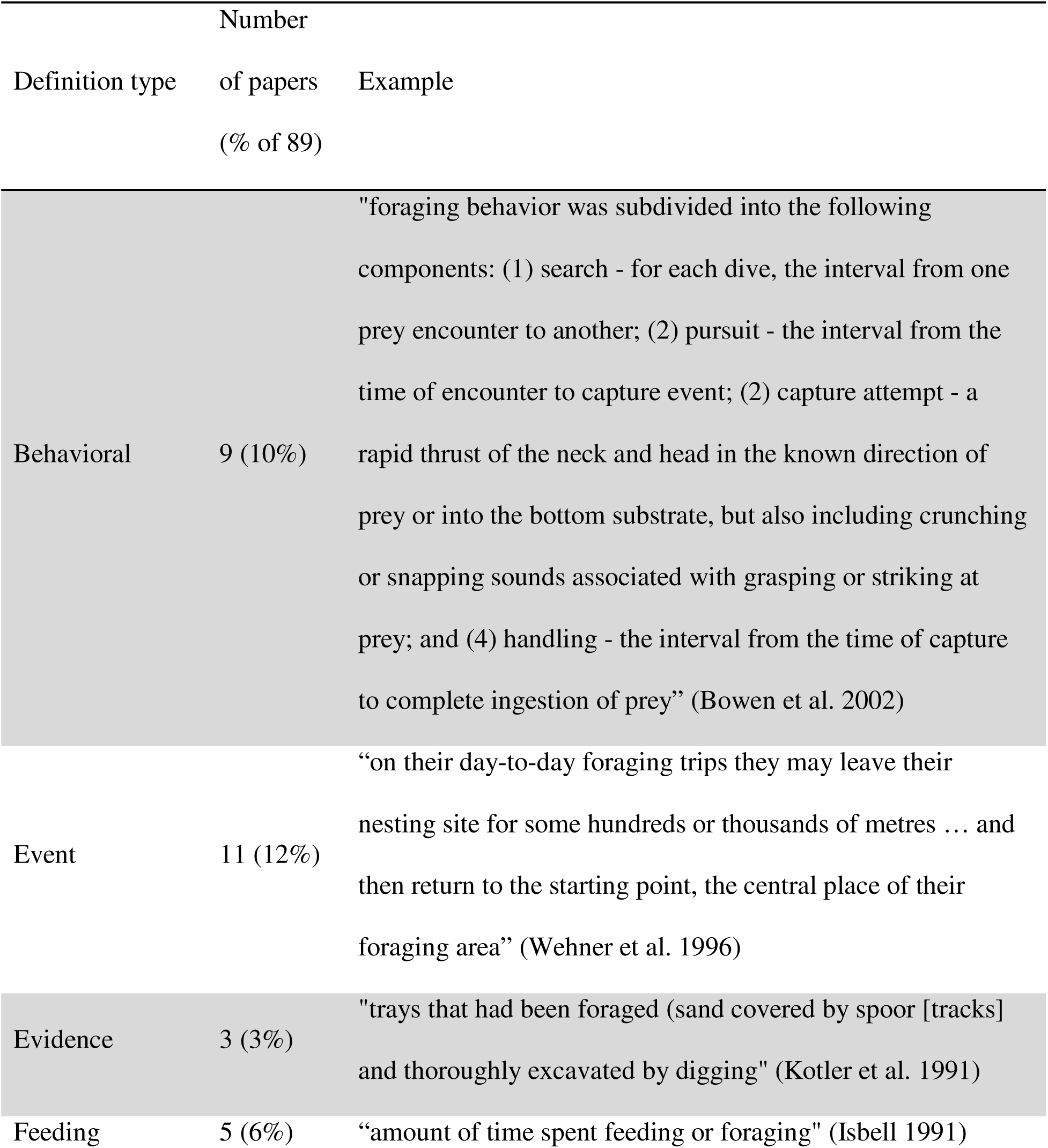
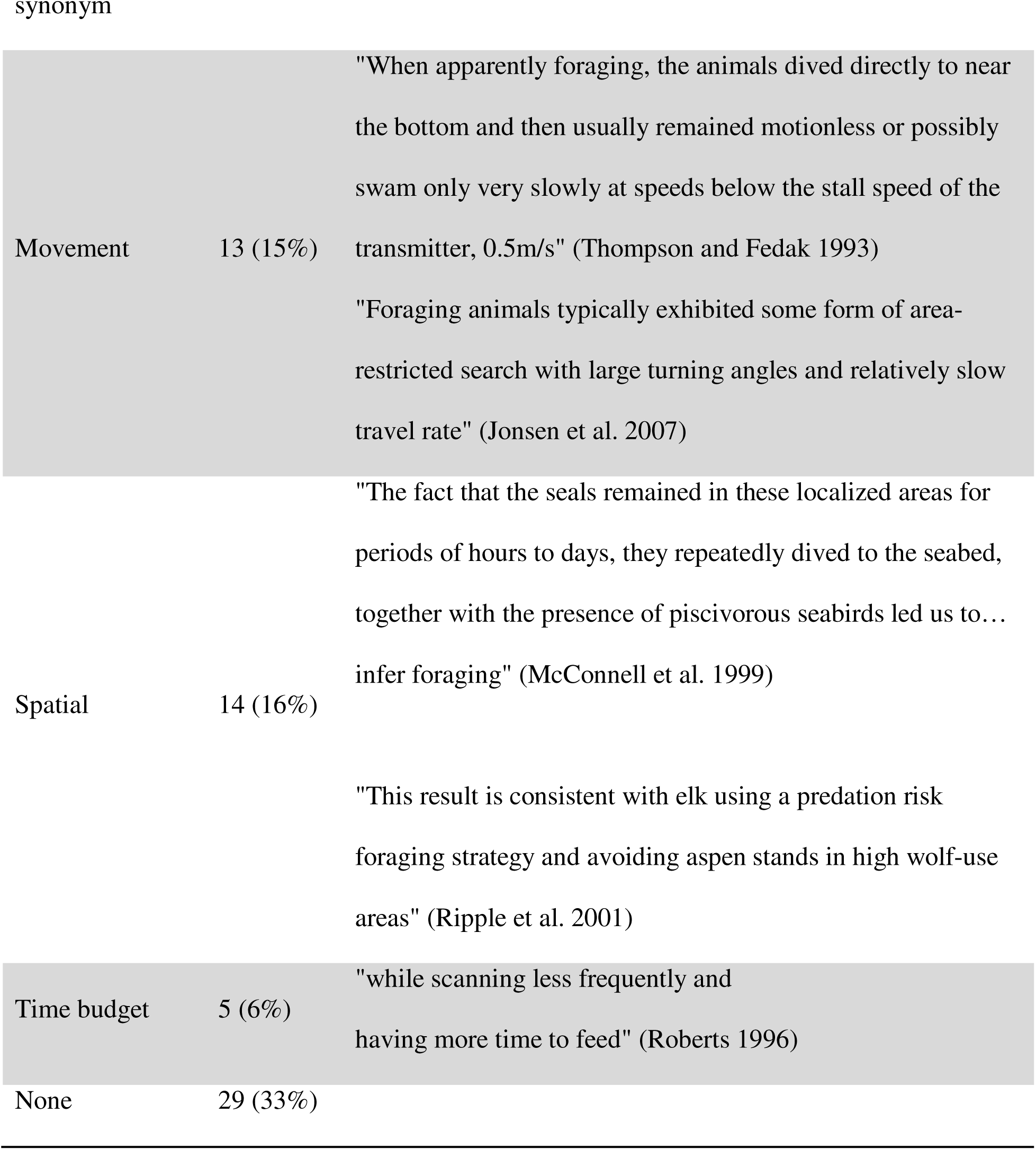
Summary of the various types of definitions of foraging included in the top 100 studies of foraging behavior using direct observational or remote approaches.

## Appendix 2: Cluster analysis and categorising approaches

Our initial search terms and categorization contrasted classical ethology and remotely sensed approaches as two distinct fields of studying foraging behavior. While the two are undoubtedly different, they are also umbrella terms that each incorporate many different technologies and processes for observing animal behavior.

The distinction between spatial data and biologgers as two disparate remote technologies was apparent from the beginning of this review. They are grouped together as remotely sensed data, but the insights offered from a GPS collar as opposed to an animal carrying an accelerometer, temperature logger, and onboard camera differ by kind and not degree. From the perspective of the information obtained on the focal animal, an accelerometer and audio or video logger is more comparable to direct observations of movement and behavior than to purely spatial data. In fact, many studies involving direct observations use video recordings of the focal animal that are later analysed and scored for the behavior of interest. For the purposes of the final data, we do not see any meaningful difference between a researcher standing in the field and writing down individual bites by a grazing ungulate, as opposed to another researcher setting up a tripod and digital camera, and later watching the video and noting bite rate from it. The intervention of the camera into these observations does not change the information obtained or the insight permitted by these two researchers. This distinction is further complicated when the full range of research methods is considered: where does one draw the line between setting up and recording an animal while you are physically present, setting up a camera and leaving it recording at a burrow or den site, deploying motion-activated camera traps and obtaining video records from free-ranging animals, or deploying a video camera on a collared animal and scoring their behavior from a first-person perspective?

We used a set of six dichotomous criteria (defined in Table A2) to describe each of 35 unique methods, and then determined the optimal grouping of these methods using *k*-means cluster analysis. While research methods might be best described along a continuum of remote observation to classical ethology, some number of discrete categories was required to compare between approaches and draw conclusions. The cluster tree for various values of *k* showed that from *k* = 5 or 6 onwards, the clusters are essentially stable and there is marginal value to including additional groups (Figure A1). Manually inspecting the clusters for *k* = 4, 5, and 6 showed that two of the clusters were consistent throughout, one comprising different ethological methods (n = 9) and one including various biologger technologies (n = 12). The remaining methods (n = 14) were split into two, three, or four groups; we decided that these two groups (*k* = 4) were too broad and lumped together quite disparate methods, whereas four groups (*k* = 6) split the data more than was necessary and created quite small groups. For the purposes of this review, we settled on five categories of approaches as the best balance of allowing for more than solely binary comparisons of approaches, while still being manageable and not overly complex. This judgement was supported by visually inspecting the scree plot of within-cluster sum of squares for each value of *k* (Figure A2). The point where this curve bends sharply towards the horizontal is thought to be the optimal value of *k* (where the curve “kinks”; Makles, 2012), and *k* = 5 more or less aligns with this point.

**Table A2.**
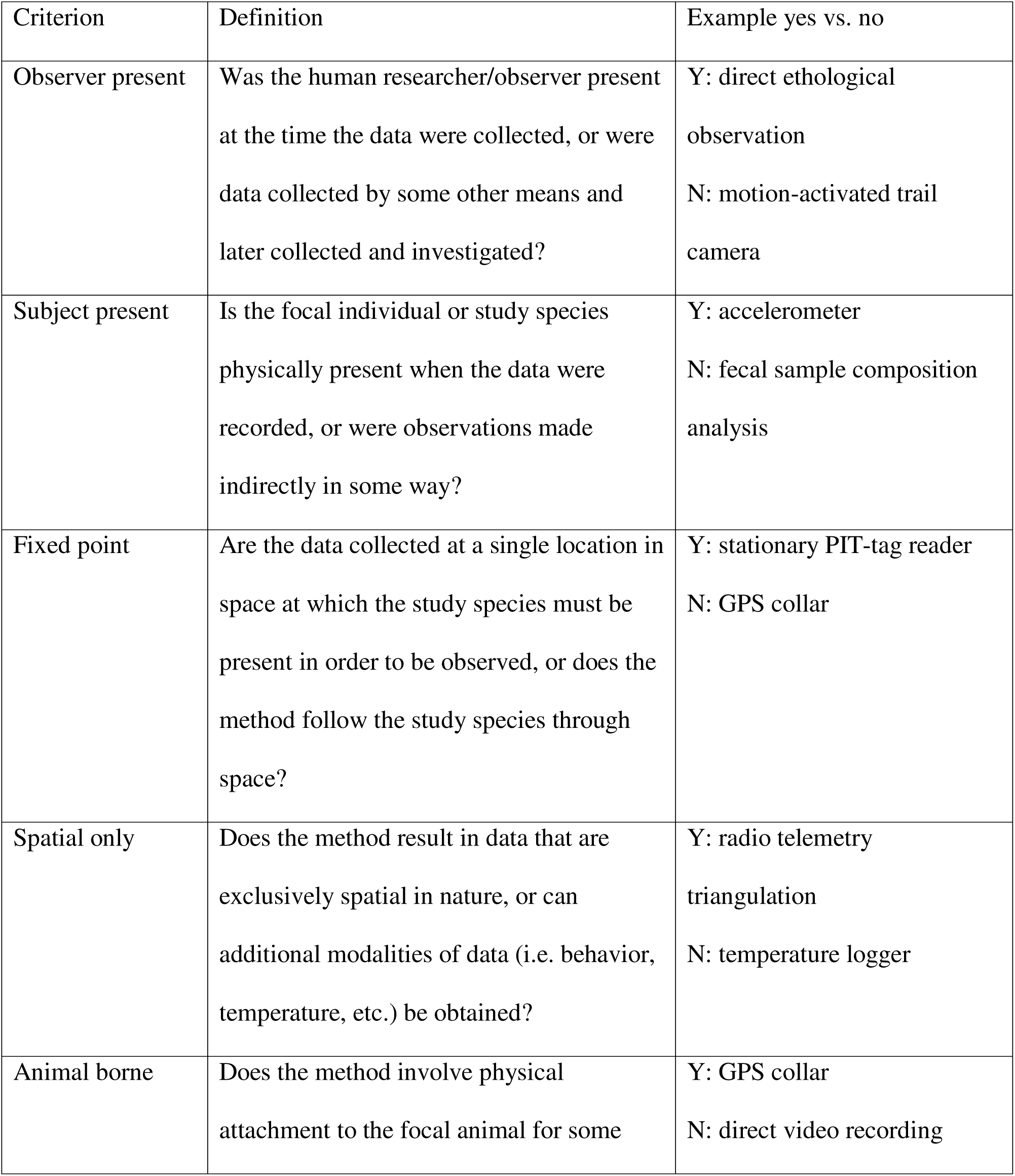
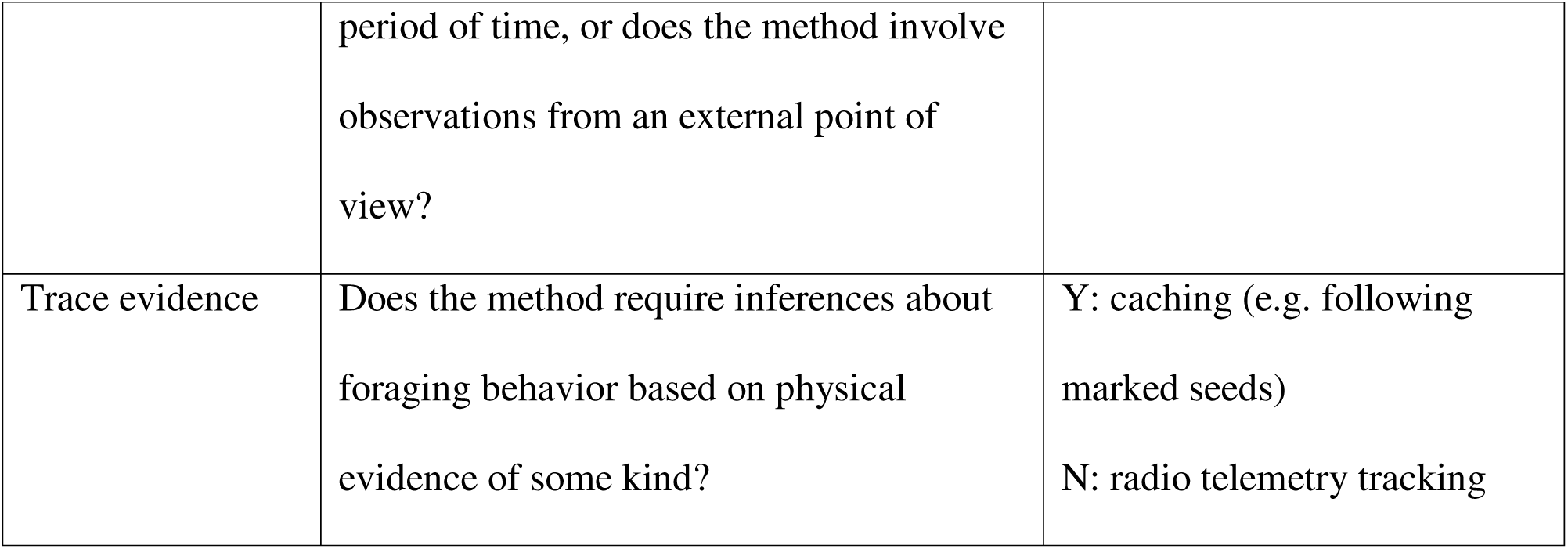
Study methods were systematically described and sorted using six dichotomous criteria, defined below.

**Figure A1.**
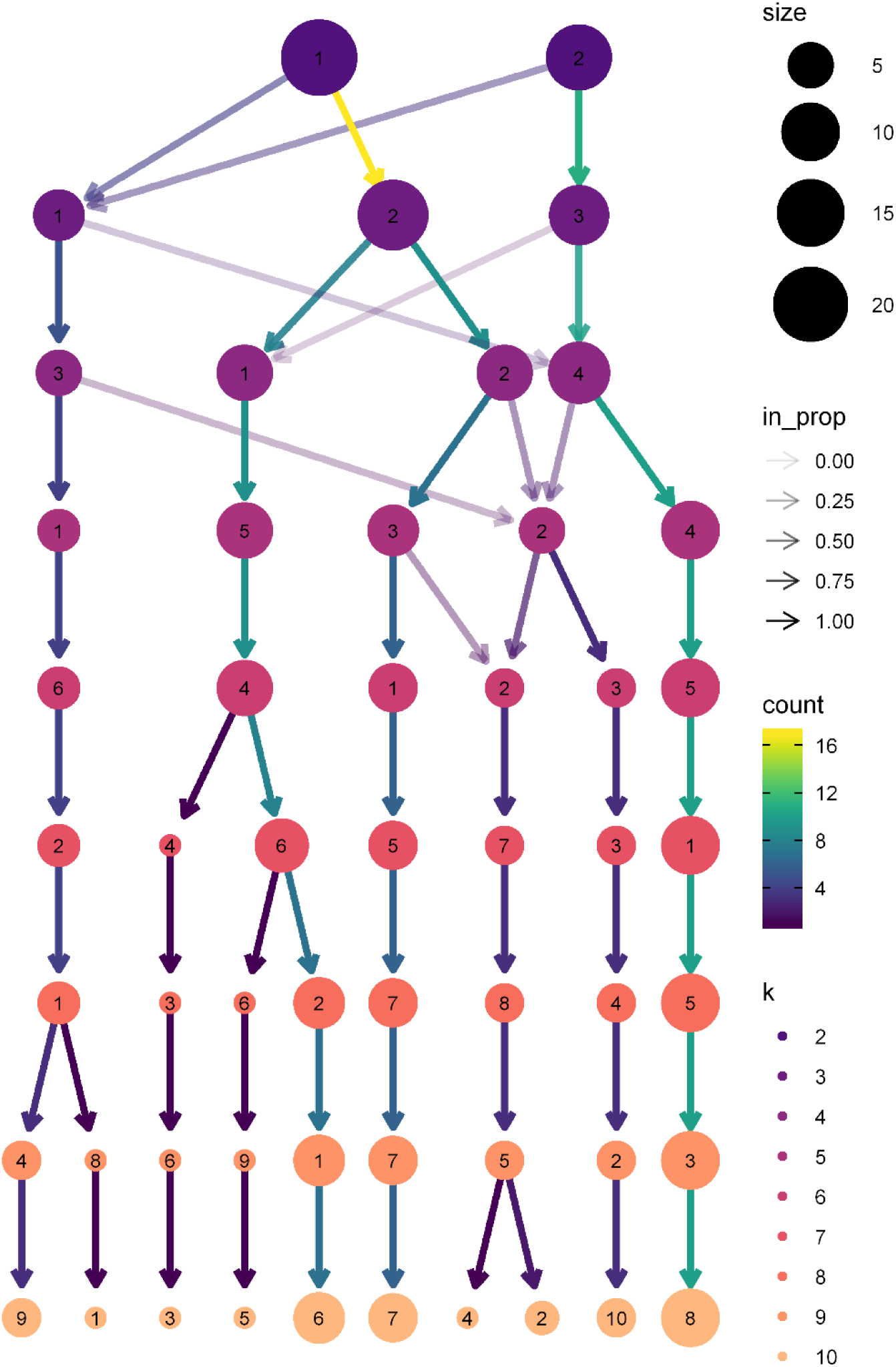
Graphical depiction of *k*-means clustering of 35 methods of studying foraging behavior for values of *k* between 2 and 10. Node color reflects each successive value of k (i.e., all nodes of the same color on the same horizontal line are clusters within one categorization scheme). Node size represents the number of methods within each cluster, and node labels refer to the cluster number within each categorization (i.e., for k = *4*, there are four clusters labelled 1, 2, 3, and 4). Arrow transparency and color indicates the proportion and number, respectively, of methods moving from one group to another as *k* increases and additional groupings are added.

**Figure A2.**
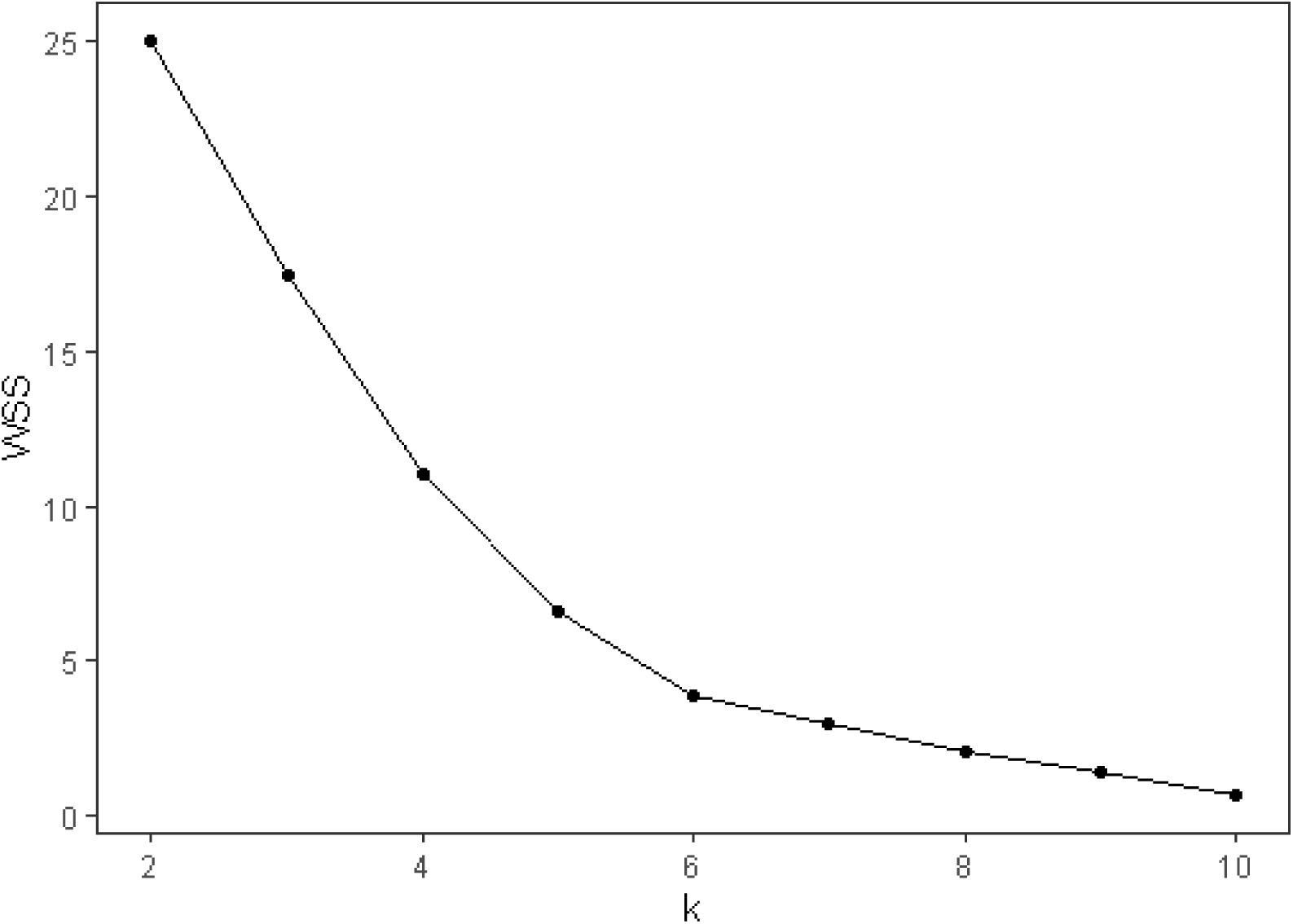
Scree plot of the total sum of squares within clusters (WSS) for each value of *k*. K-means cluster analysis aims to minimize the sum of squares within clusters without dividing the data unnecessarily, such that the optimal value of k is where the curve of WSS kinks or bends sharply (Makles 2012). In this analysis, the curve bends fairly smoothly without any obvious vertex, but *k* = 5 or 6 approximates this point.

## Notes

https://github.com/jghendrix/foraging-tech/

### Competing Interest Statement

The authors have declared no competing interest.

